# Resolving the time course of visual and auditory object categorization

**DOI:** 10.1101/2021.11.25.470008

**Authors:** Polina Iamshchinina, Agnessa Karapetian, Daniel Kaiser, Radoslaw M. Cichy

**Affiliations:** Department of Education and Psychology, Freie Universität Berlin, Habelschwerdter Allee 45, Berlin 14195, Germany; Berlin School of Mind and Brain, Humboldt-Universität zu Berlin, Unter den Linden 6, Berlin 10099, Germany; Mathematical Institute, Department of Mathematics and Computer Science, Physics, Geography, Justus-Liebig-Universität Gießen, Arndtstraße 2, 35392 Gießen; Center for Mind, Brain and Behavior (CMBB), Philipps-Universität Marburg and Justus-Liebig-Universität Gießen, Hans-Meerwein-Straße 6, 35032 Marburg, Germany

**Author notes:** These authors contributed equally (Daniel Kaiser, Radoslaw M. Cichy).

## Abstract

Humans can effortlessly categorize objects, both when they are conveyed through visual images and spoken words. To resolve the neural correlates of object categorization, studies have so far primarily focused on the visual modality. It is therefore still unclear how the brain extracts categorical information from auditory signals. In the current study we used EEG (N=47) and time-resolved multivariate pattern analysis to investigate (1) the time course with which object category information emerges in the auditory modality and (2) how the representational transition from individual object identification to category representation compares between the auditory modality and the visual modality. Our results show that (1) that auditory object category representations can be reliably extracted from EEG signals and (2) a similar representational transition occurs in the visual and auditory modalities, where an initial representation at the individual-object level is followed by a subsequent representation of the objects‘ category membership. Altogether, our results suggest an analogous hierarchy of information processing across sensory channels. However, we did not find evidence for a shared supra-modal code, suggesting that the contents of the different sensory hierarchies are ultimately modality-unique.

## 1. Introduction

Whether we see a pineapple or hear somebody say “pineapple”, we can rapidly and effortlessly infer key properties of the object: for instance, we can confidently say that a pineapple is a natural, inanimate object. Such categorization processes are essential for utilizing object knowledge in an efficient way. So far, the studies in the field of object recognition have been investigating the neural correlates of object categorization primarily in the visual modality. Using fMRI and M/EEG, researchers have identified a gradual progression from visual representations of individual objects to more abstract representations of an object’s category, both along the ventral visual hierarchy and across processing time (Shinkareva et al., 2011; Fairhall & Caramazza, 2013; Cichy et al., 2014; Proklova et al., 2016; Kaiser, Azzalini & Peelen, 2016; Kumar et al., 2017; Leonardelli, Fait & Fairhall, 2019).

By contrast, studies seldom focus on object categorization from auditory inputs such as linguistic utterances. Few fMRI studies have pinpointed categorical coding for auditory stimuli to superior temporal and medial frontal cortex (Simanova et al., 2012; Jung, Fait & Walther, 2018). Only one EEG study so far has tried to systematically compare the time course of visual and auditory abstract information but did not succeed in reliably establishing category information for auditory stimuli (Simanova et al., 2010). A different line of research investigates the time course of semantic word analysis using event-related potentials: when participants read or listen to full sentences, an N400 ERP component is observed in response to a categorical misattribution of words (Fischler et al., 1984; Fischler, 1985; Núñez-Peña & Honrubia-Serrano, 2005; Hnazaee & Hulle, 2017). Yet, the N400 was found to coincide with a wide spectrum of semantic incongruencies (Kutas & Federmeier, 2011) and it is currently unclear to what extent the waveform is specific to categorization processes.

Here we pose two critical questions about object recognition from auditory inputs. First, how does object category information dynamically emerge from auditory inputs? Second, is there a representational transition from individual object identification to category membership attribution in the auditory modality and how does it qualitatively compare to the dynamics of object categorization in the visual modality?

To answer these questions, we tracked the emergence of visual and auditory category information in EEG signals. We used a paradigm commonly used in studies of visual object recognition: To evoke automatic category processing and to avoid any context- or task-driven modulations, we presented participants (N=48) with images of objects and spoken words corresponding to the same objects while they were doing an orthogonal 1-back task. Objects belonged to three category dimensions, based on object animacy, size, and movement, which were previously shown to explain substantial variance in object representations (Huth et al., 2012). We used time-resolved multivariate pattern analysis (MVPA) on the resulting EEG data to identify the temporal transition from object-specific to category-defining representations. First, we found that EEG responses after 300 ms of processing form a neural correlate of object categorization in the auditory modality. Second, by tracking representations of individual objects and categories across time, we demonstrate that sensory signals similarly traverse the stages of object identification and categorization in both modalities, suggesting that the perceptual hierarchy established in vision is qualitatively similar for other sensory channels.

## 2. Materials and methods

### 2.1. Participants

51 healthy adult participants took part in the study. Four participants had to be excluded due to excessive noise in the data, so that the final sample consisted of 48 participants (mean age ± std = 25.02 ± 5.04; 33 female). The study was conducted at the Center for Cognitive Neuroscience Berlin. Participants were compensated with credits or a monetary reward. All participants were native German speakers with normal or corrected-to-normal vision. All participants provided informed written consent. The study was approved by the ethics committee of the Department of Education and Psychology at Freie Universität Berlin.

### 2.2. Stimuli

The stimulus set was composed of 48 objects each presented as images or as spoken words in German (Figure 2A). The objects were organized according to three orthogonal dimensions, each divided in two categorical divisions: size (big or small), movement (moving or non-moving) and naturalness (natural or man-made (artificial)). Each item was assigned to one unique combination of categories along these dimensions (e.g., a baby is small, moving and natural). The stimulus set was balanced such that each categorical division included one half of the stimulus set (24 objects). The choice of categorical divisions was based on previous studies demonstrating that the semantic dimensions spanning these categories yield reliable neural representations independent of experimental design or neuroimaging method (Huth et al., 2012; Cichy et al., 2014; Carlson et al., 2013). The images were selected from Google images using a copyright-free search filter. The size of the images was 400 × 400 pixels. Recordings of the words being spoken were made by the investigators. The words were matched for word length (mean length ± std = 6.93 ± 2.09 letters; mean number of syllables ± std = 2.5 ± 0.51).

### 2.3 Experimental procedure

The experiment was divided into auditory and visual runs. It always started with 8 auditory runs, followed by a short break and 6 visual runs. The auditory runs were always first to prevent participants from imagining the exact same object they had seen during the visual runs and therefore to avoid possible contamination of the results of crossmodal decoding with visual mental imagery during auditory word presentation. We included two more auditory runs than visual runs, as based on pilot data we expected lower signal-to-noise ratio for auditory signals. Each run consisted of 300 trials and lasted 6 minutes. Each stimulus was repeated 5 times per run, thus each stimulus was presented 40 times over the auditory runs and 30 times over the visual runs.

In visual trials, a pseudo-randomly selected stimulus was presented on a gray screen at a visual angle of 4.24°, overlaid with a black fixation cross. In auditory trials, only the fixation cross was present while participants heard the words. For both modalities, stimulus presentation was preceded by a frame with a red fixation cross to aid attention preparation. On 20% of trials the stimulus was repeated and participants were tasked to press a button (Figure 1B). These one-back repetition trials were excluded from the analysis. To match stimulus durations across modalities, we created a distribution of durations for the visual stimuli based on the duration of the auditory stimuli (mean duration ± std = 690 ms ± 176 ms) and randomly assigned these durations to visual stimuli. The inter-trial interval (ITI) was jittered (500 ± 50 ms). The ITI after one-back repetition trials was 200 ms longer to allow enough time for a button press.

**Figure 1.**
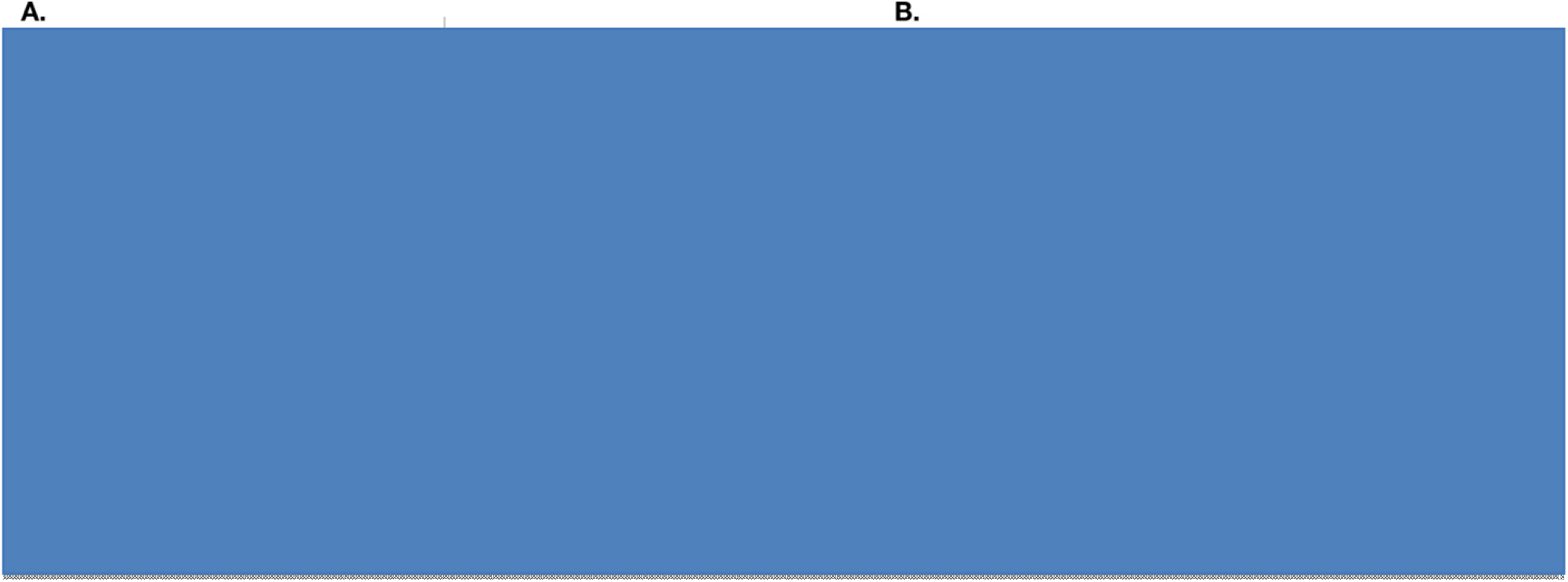
Experimental design. **A.** The stimulus set consisted of 48 objects belonging to 3 categorical divisions. In the visual runs, participants viewed images of these objects, while in the auditory runs, they heard the names of the objects. **B.** Both in visual (left) and auditory (right) runs, participants were presented with a random sequence of stimuli. Their task was to press a button when two subsequent stimuli were identical (one-back task).

### 2.4 EEG recording

EEG data were collected using the Easycap 64-electrode system and BrainVision Recorder. The participants wore actiCAP elastic caps, connected to 64 active scalp electrodes: 63 EEG electrodes and one reference electrode (Fz). The activity was amplified using the actiCHamp amplifier, sampled at 1000 Hz, and filtered online between 0.5 and 70 Hz.

### 2.5 Data preprocessing

The data were preprocessed offline using the FieldTrip toolbox (Oostenveld et al., 2011) for MATLAB (2018b). The data were first segmented into epochs from 200 ms before stimulus onset to 800 ms post-stimulus. Afterwards, the data were downsampled to 200 Hz and trials with artifacts were removed (i.e, a trial is excluded if standardized deviations from the mean of all channels in it are larger than 20, for details see jump artifact in fieldtrip toolbox). Noisy channels and trials were removed based on visual inspection.

### 2.6 Classification analysis

Multivariate pattern analysis (MVPA) was carried out using linear support vector machines (SVMs; libsvm: http://www.csie.ntu.edu.tw/~cjlin/libsvm/) with a fixed cost parameter (c=1). We performed separate classification analyses on electrode patterns from every millisecond of the trial epoch. We performed classification on the object- and on the category-level, as explained in the following.

For object-level classification, all the trials were sorted according to the object presented in each particular trial (e.g., ballerina or banana). Within every object condition, we randomly assigned trials into three groups, and then averaged the trials within each of these groups to form three “super-trials”. The super-trials were then normalized using multivariate noise normalization (Guggenmos et al., 2018) to downscale channels with high noise covariance and thereby improve signal reliability.

The resulting data were used to perform pairwise classification between all possible pairs of objects. Specifically, we trained a classifier with a 3-fold cross-validation approach using 2 out of the 3 super-trials from each of the 2 object conditions (ballerina vs. banana). We tested the classifier on the left-out super-trial. This classification procedure was repeated 100 times, with different random assignments of trials into the 3 super-trials. Classification accuracies were averaged across these 100 repetitions. Finally, by averaging all pairwise classification accuracies, we obtained a measure of object-level classification.

For category-level classification, all the trials were sorted according to the object presented in each particular trial and then averaged for each object. Then the object-level averages were sorted by category: this was done three separate times, for each of the three category dimensions (e.g., moving vs. non-moving, Figure 1A). Within each category division (e.g., moving objects), we randomly assigned the object averages into three groups, and then averaged within each of these groups to form three “super-trials”. Classification was performed in a leave-one-out scheme across the 3 super-trials as outlined above. Critically, the initial averaging of trials at the object level prevented classifiers from training and testing on trials with the same object, thereby probing category-level representations independent of the low-level properties of individual objects in our stimulus set. We again repeated the classification procedure 100 times, with different assignments of the object-level averages into super-trials and averaged the decoding accuracies across these repetitions. Finally, by averaging across all three category distinctions, we obtained a measure of category-level classification.

### 2.7 Statistical analysis

We used non-parametric statistical inference (Nichols & Holmes, 2002; Maris & Oostenveld, 2007), which does not make assumptions about the distribution of the data. Permutation tests were used for cluster-size inference, in which we randomly multiplied the participant-specific data (e.g., EEG decoding accuracies) with +1 or −1 for 10,000 times to create a null distribution. All tests were one-sided against a 50% chance level and thresholded at p-value < 0.05.

We used a non-parametric test to calculate differences in the onsets of significant decoding between two conditions (object/category information), that is, the difference between the time points at which objects were first discriminated. To estimate if the decoding reaches significance in one condition reliably earlier/later than in the other condition, we created 1,000 bootstrapped samples by sampling the participant-specific data with replacement and performing a sign-permutation test at each iteration. Thus, every iteration yielded an onset of significant decoding. Then, combining the onsets from all the iterations yielded an empirical distribution of onsets of significant decoding in two conditions of interest. Then, we subtracted the onsets estimated in one condition from the onsets estimated in the other condition (object information – category information). We calculated p-values (one-tail) by dividing the number of bootstrapped samples with differences greater than 0 (e.g., those samples in which the onset of object information is early than the onset of category information) by the overall number of samples (1,000).

## 3. Results

### 3.1 The time course of visual object representations

Based on previous studies (Shinkareva et al., 2011; Fairhall & Caramazza, 2013; Cichy et al., 2014, Kumar et al., 2017; Leornardelli, Fait & Fairhall, 2019) revealing a processing hierarchy starting from visual object representations to more abstract category representations, we expected that we could uncover both types of representations from the EEG signals evoked by the object images. Further, we expected that object-level representations would emerge earlier than category representations.

We found that EEG signals conveyed significant visual object information from 80 ms to 800 ms after image onset (Figure 2A), and significant category information from 135 ms to 800 ms (Figure 2B). Notably, category information only emerged significantly after object information (test for onset difference: p=0.001, see Methods), revealing a temporal progression from visual to more abstract representations. Note that given the differences in the two decoding approaches (see Methods), absolute decoding accuracies are not directly comparable for the two analyses.

**Figure 2.**
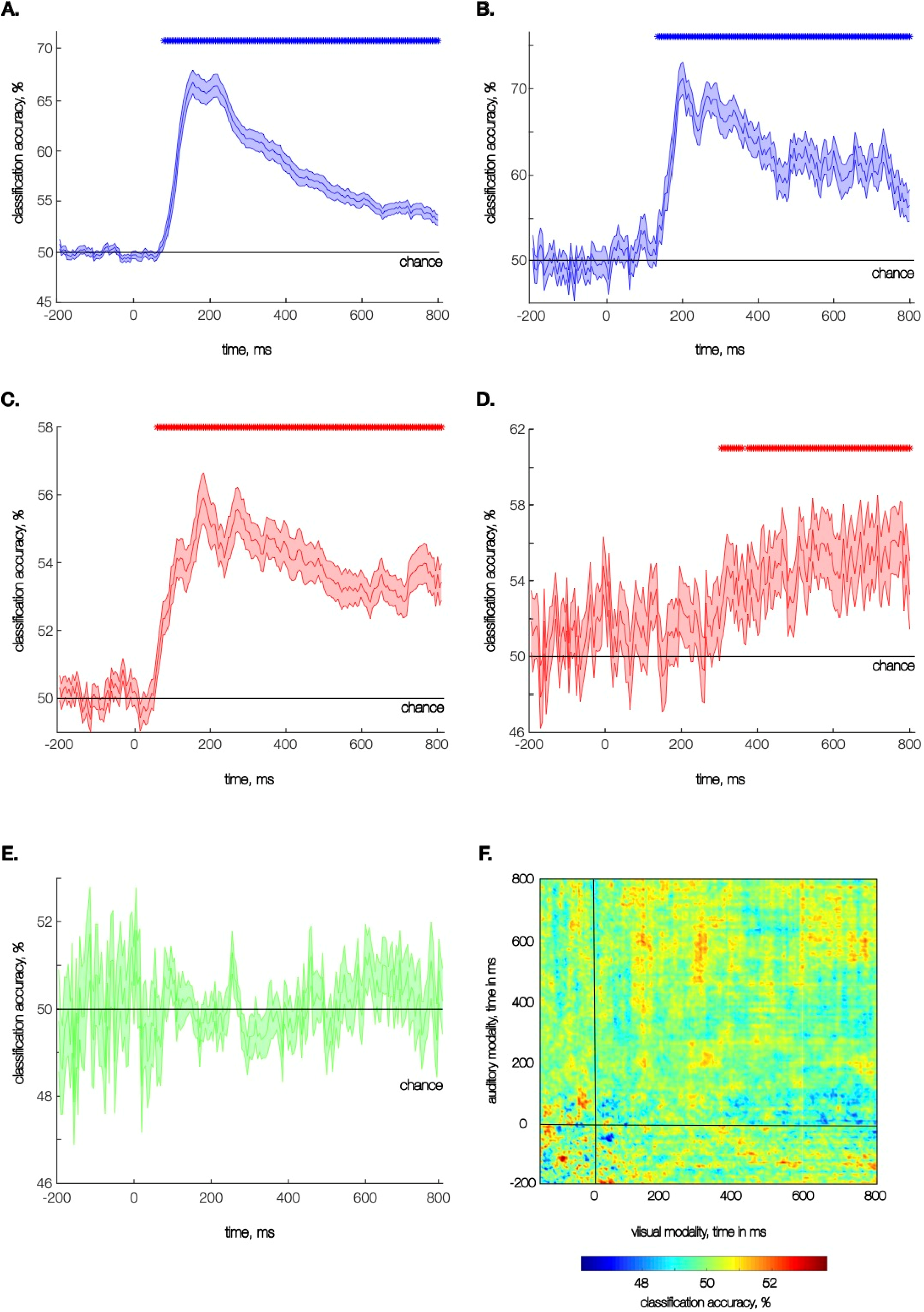
Classification Results. **A.** Object information time course in the visual modality **B.** Category information time course in the visual modality averaged across decoding results obtained for each pair of categorical divisions. **C.** Object information time course in the auditory modality **D.** Category information time course in the auditory modality averaged across decoding results obtained for each pair of categorical divisions. **E.** Category information time course, where classifiers were trained on one modality and tested on the other modality. Results are averaged for both train/test directions. **F.** Time generalization results for category information, where classifiers were trained on one modality and tested on the other modality. Results are averaged for both train/test directions. The onset of the stimulus presentation is at 0 ms. Rows of asterisks indicate significant time points (p<0.05, corrected for multiple comparisons). Error bars denote between-participant SEM.

### 3.2 The time course of auditory object representations

Next, we tested whether we could also retrieve object category information when the objects were conveyed through the auditory modality and whether in this case a similar progression from object-level representations to category representations can be observed.

As for the visual modality, we found temporally sustained object information from 60 ms to 800 ms after word onset (Figure 2C). We also found significant category information from 305 ms to 800 ms (Figure 2D), showcasing that object category can be reliably retrieved from auditory brain signals. Further, this category information emerged significantly after object-level information (p=0.038), suggesting a similar representational transition towards more abstract, categorical stages of processing in the visual and auditory modalities.

### 3.3 Commonalities between visual and auditory representations

Finally, we asked whether categorical object representations present in both modalities reflect a convergence towards conceptual representations that are modality-independent. In case of such a convergence, we should be able to cross-classify object category across visual and auditory brain signals. For cross-classification, we trained a classifier on response patterns to each pair of conditions in one modality and tested the classifier on response patterns to the same pairs of conditions from the other modality. In this analysis, no significant cross-decoding was found at any time point across the epoch (Figure 2E).

However, the temporal processing cascades do not necessarily need to match between the visual and auditory modalities. We therefore also performed a time generalization analysis, in which we trained classifiers on each time point in one modality and tested them on all timepoints in the other modality. Also here, we found no significant cross-decoding. These results indicate that despite the robust category information in both modalities, there is no shared conceptual code for object representation detectable on the level of scalp electrode patterns in our data (Figure 2F).

## 4. Discussion

In this study, we investigated the temporal dynamics of object category processing in the visual and auditory modalities. Specifically, we were interested to know when object category information emerges in the auditory modality and whether the representational transition from object-to category level in auditory modality is qualitatively similar to that in vision. Our results show that auditory object category representations can be reliably extracted from EEG signals. Further, they show that there is an analogous representational transition in the visual and auditory modalities, with an initial representation at the individual-object level, and a subsequent representation of the objects‘ category membership.

This representational transition has been firmly established in the visual domain before (e.g., Cichy et al., 2014; Kumar et al., 2017). Crucially, our study also demonstrates the temporal dynamics of auditory object representations at these different levels of abstraction. These results extend previous fMRI research (Simanova et al., 2012; Jung, Fait & Walther, 2018) that showed categorical information arising from auditory inputs in superior temporal and medial frontal gyri: Our findings suggest that these categorical representations emerge only well after object-level representations, from around 300 ms after the word onset. The time course of category representation obtained in our study corresponds to the one previously obtained for written words (Leonardelli et al., 2018), pointing at similarities in processing visual and auditory language information.

The auditory categorical signals in our study temporally align with the occurrence of the N400 component elicited in response to semantically incongruent spoken words (300-900 ms, Bentin, Kutas & Hillyard, 1993). Several studies specifically demonstrated that the N400 can be evoked by a categorical misattribution of a word (Fischler et al., 1984; Fischler, 1985; Fujihara et al., 1998; Núñez-Peña & Honrubia-Serrano, 2005; Hnazaee & Hulle, 2017), thereby hinting at the component as a specific timestamp for word categorization. Building on this previous research, our findings suggest that extracting the categorical membership during spoken word perception may partially underlie the emergence of N400 in response to categorical misattribution. Further investigation is needed to establish the role of category discrimination in the process of word meaning extraction (Kocagoncu et al., 2017; Friederici, 2002).

Although we found robust category information in both the visual and auditory modalities, we did not find evidence for a transformation of representations from modality-specific codes to modality-independent conceptual representations, as evidenced by the absence of significant cross-modal decoding. In contrast, two previous fMRI studies identified representations that generalize across the auditory and visual modalities in inferior temporal, inferior frontal, and middle frontal cortices (Simanova et al., 2012; Jung, Fait & Walther, 2018). Why did we not find evidence for such representations here? First, cross-modal convergence of representations may be particular to visual and linguistic information being conveyed through the same modality, as for instance for images and written words (Simanova et al., 2012). Second, the current study used an orthogonal task to measure the process of automatic category extraction, which might not sufficiently engage late, modality-independent processes. Future studies could employ tasks, such as category verification or story listening/reading (Huth et al., 2016; Deniz et al., 2019; Popham et al., 2021) that encourage deep processing of words and their context in modality-independent rather than modality-focused details. Third, we cannot exclude the possibility that M/EEG scalp sensor patterns lack the sensitivity to uncover the subtle signal differences essential for the readout of modality-unspecifc contents (Giari et al., 2020), while such differences can be revealed with spatially precise fMRI recording (Proklova et al., 2016, 2019).

Together, our results elucidate the time course of categorical object coding in the visual and auditory modalities. Further, they establish commonalities in the representational transition from object-level information to categorical representations across the two modalities, suggesting a similarity in the hierarchy of information processing across sensory channels.

